# *In Vitro* Validation of Computationally Predicted Oncogenic Driver Mutations in EGFR Tyrosine Kinase Domain

**DOI:** 10.1101/2023.01.29.526080

**Authors:** Vidya P Warrier, Anoosha Paruchuri, Sakthivel Ramasamy, Michael M Gromiha, Devarajan Karunagaran

## Abstract

Epidermal Growth Factor Receptor (EGFR) signaling is known to play essential roles in growth and development; nevertheless, overexpression and mutation of EGFR have been reported in several cancers. Non-small cell lung cancer (NSCLC), the most observed type of lung cancer, harbors the highest number of EGFR tyrosine kinase mutations and therefore, EGFR has become an important therapeutic target for treatment of these tumors. Tyrosine Kinase Inhibitors (TKIs) are found to be effective in patients whose tumors contain activating mutations in the tyrosine kinase region of the receptor. This would seem to be beneficial in the treatment of EGFR mutation-positive NSCLC patients but the activating mutations should be sensitive to TKIs. Earlier, a machine learning approach was developed to classify single amino acid polymorphisms (SAPs) in EGFR into driver (cancer-causing) and passenger (neutral) mutations using structural and functional features (Anoosha *et al*., 2015). This study screened all possible point mutations in EGFR and predicted a list of mutations with high probability of being a driver or a passenger. From this list, we selected 2 mutations (G729E and G719F) with high evolutionary conservation score for *in vitro* validation. If proven to be oncogenic drivers and sensitive to EGFR TKIs, these mutations can aid in the early diagnosis and successful therapy of EGFR mutation-positive NSCLC.

## Introduction

Lung cancer, a malignant tumor characterized by uncontrolled cell growth in lung tissue, is the leading cause of cancer-related deaths among men and ranks second next to breast cancer among women worldwide (GLOBOCAN 2018). Even though smoking and tobacco usage are the major risk factors involved in 90% of lung cancer cases, environmental risk factors (such as radon gas, asbestos, air pollution, and chronic infections) as well as multiple inherited and acquired aberrations in oncogenic or tumor suppressor signaling pathways are also involved. Aberrations in Epidermal Growth Factor Receptor (EGFR) signaling are common in lung cancer. EGFR is a transmembrane receptor tyrosine kinase protein that is activated by ligands such as Epidermal Growth Factor (EGF) and initiates many signal transduction cascades like MAPK, PI3K-Akt, STAT5, and PLCγ-PKC pathways leading to cell proliferation and development (Yarden and Schlessinger, 1987). Overexpression and mutation of EGFR lead to constitutive activation and aberrant signaling in breast, lung, colon, stomach, pancreatic, head and neck, ovarian, kidney, prostate cancer, and gliomas. Somatic mutations of EGFR dominate (approximately 26% according to COSMIC database) as compared to other common gene mutations (TP53 and KRAS) in lung cancer. Although, mutations of EGFR occur in the extracellular region, tyrosine kinase region and the transmembrane region, tyrosine kinase mutations are predominant in lung cancer.

The therapeutic strategies to inhibit EGFR fall into two major categories: monoclonal antibodies (mAbs) that target the extracellular region of EGFR and Tyrosine kinase inhibitors (TKIs) that target the tyrosine kinase region. The TKIs can further be classified as first generation TKIs (gefitinib, erlotinib), second generation TKIs (afatinib, dacomitinib), and third generation TKIs (osimertinib). All these TKIs work by competing with ATP molecule for the ATP-binding cleft of EGFR. The first-generation inhibitors are reversible while the second and third generation inhibitors are irreversible in nature. Interestingly, gefitinib and erlotinib are found to be more active in a subset of patients with activating mutations in the exons 18-21, that surround the ATP cleft and TKI binding pocket of EGFR. Mutations occurring in this area may reposition the critical residues, thereby stabilizing their interactions with both ATP and the TKI molecule (Lynch *et al*., 2004). Analysis of mutations would therefore provide more insight into mechanisms that regulate EGFR activation and help to design potent inhibitors targeting the mutant receptor.

Earlier, a machine learning approach was developed in our lab to classify single amino acid polymorphisms (SAPs) in EGFR into driver (cancer-causing) and passenger (neutral) mutations using structural and functional features (Anoosha *et al*., 2015). Our study screened all possible point mutations in EGFR and predicted a list of mutations with high probability of being a driver or a passenger. From this list, we selected 2 mutations that gave a high evolutionary conservation score (Consurf analysis data) for our study: G729E and G719F. Interestingly, G729E was reported in 2 lung cancer samples and 1 skin cancer sample (Yi *et al*., 2014, Payne *et al*., 2006, Penzel *et al*., 2011) but so far, no further study has been conducted to validate other predicted mutations in EGFR. Hence, the overall idea of the proposed study is the functional analysis of the 2 predicted mutations in EGFR and to check whether the computationally predicted driver mutations can be confirmed *in vitro*.

## Materials and Methods

### Cell Lines and Culture

The cell lines NIH-3T3, HEK-293T and A549 chosen for the study were obtained from ATCC. NIH-3T3 is a mouse fibroblast cell line with minimal level of endogenous EGFR expression while A549 is a human lung NSCLC cell line that overexpresses EGFR and this cell line was used as a positive control in our study. HEK-293, a human embryonic kidney cell line was used for retroviral transduction to establish stable cell lines. All cell lines were cultured in Dulbecco’s Modified Eagle’s Medium (DMEM, Gibco, Thermoscientific) containing 10% Fetal Bovine Serum (FBS, Gibco, Thermoscientific) and antibiotics (100 U/mL penicillin and 100 μg/mL streptomycin) in a humidified atmosphere of 5% CO2 at 37°C.

### Site-Directed Mutagenesis

EGFR wild type (pBABE-puro EGFR WT), EGFR known driver mutant L858R construct (pBABE-puro EGFR L858R), and empty vector (pBABE-puro) were purchased from Addgene. The wild type and mutant constructs were confirmed by sequencing. The mutation specific primers 5’-GGCACGGTGTATAAGGAACTCTGGATCCCAGAA-3’ (G729E forward), 5’-TTCTGGGATCCAGAGTTCCTTATACACCGTGCC-3’(G729reverse), 5’ -CAGATTTTGGGCGGGCCAAACTGCT-3’ (G719Fforward) and 5’-AGCAGTTTGGCCCGCCCAAAATCTG-3’ (G719F reverse) for site-directed mutagenesis were designed and purchased from Eurofins. The point mutations G729E and G719F were introduced into the EGFR-WT expression vector by PCR using Phusion high fidelity DNA polymerase (New England Biolabs). PCR product obtained was digested by methylation-specific enzyme *DpnI* (New England Biolabs) and transformed into bacteria for obtaining isolated colonies. Further, plasmids were isolated from the primary culture and the presence or absence of mutations was verified by sequencing.

### Establishment of stable cell lines

Retroviruses were produced by the co-transfection of pBABE-puro EGFR constructs with pCL-Eco (packaging vector) and pCMV-VSVG (pseudotyping envelope vector) into HEK-293T cells. Target cells (NIH-3T3) were infected with these retroviruses in the presence of 8μg/mL polybrene (Santa Cruz Biotechnology). After 2 days of infection, cells were selected using puromycin (Sigma Aldrich). The minimum puromycin concentration that killed all non-transduced cells in 3-7 days was taken as the optimum concentration of puromycin for selection during transduction.

### Western Blotting

Total cell lysates (with or without 10ng/mL EGF) were prepared using RIPA lysis buffer [Tris, NaCl, EDTA, β-glycerophosphate, Triton X-100, sodium pyrophosphate, sodium orthovanadate, sodium deoxycholate, PMSF, sodium fluoride and protease inhibitor cocktail] and proteins were quantified using Bradford’s method (Bradford, 1976). Protein samples were resolved on SDS-PAGE and transferred to a PVDF membrane (Bio-Rad) using a semi-dry transfer apparatus (Orange Biosciences). Primary antibodies such as anti-EGFR, anti-phospho-EGFR, anti-AKT, anti-phospho-AKT, anti-STAT5, anti-phospho-STAT5, anti-ERK1/2, anti-phospho-ERK1/2 and the appropriate secondary antibodies were purchased from Cell Signaling or Santa Cruz Biotechnology. The protein bands were visualized using the Enhanced Chemiluminescence Kit (Bio-Rad) and detected using Chemi-Doc (Bio-Rad). The intensity of the bands was analysed by densitometry using ImageLab software (Bio-Rad).

### Soft agar anchorage independent growth assay

The cells transduced with wild type or mutant EGFR were suspended with or without EGF (10ng/mL) in 0.3% agar and complete medium (DMEM supplemented with 10% FBS) and plated onto 0.6% agar coated 6 well plates. The cells, growing on an anchorage free platform, were allowed to form colonies over a period of 7-14 days with periodical renewal of DMEM-FBS. After incubation, medium was aspirated and cells were washed with PBS, fixed with methanol, and stained with crystal violet. The colonies formed were quantified using ImageJ software (Yu *et al*., 2017).

## Results and Discussion

### Generation of EGFR G729E and EGFR G719F mutations

The mutations G729E and G719F were introduced into pBABE-puro EGFR WT plasmid construct by site-directed mutagenesis using mutation specific primers. The presence of these mutations was confirmed by sequencing.

### Optimization of puromycin concentration for stable cell line generation

The optimal concentration of puromycin for selection of transduced cells was determined by titration of non-transduced target cells against varying concentrations of puromycin for a period of 3-7 days. The optimal concentration of puromycin for selection of NIH-3T3 transduced cells was determined to be 1.50 μg/mL.

### Analysis of EGFR activity (phosphorylation status) of NIH-3T3 cells transduced with pBABE-puro construct

NIH-3T3 cells transduced with EGFR L858R expressed high levels of phosphorylated EGFR compared to cells transduced with WT EGFR as expected since L858R is a known oncogenic driver mutation (Fig.3.A). As evident in Fig.3.C, NIH-3T3 cells transduced with EGFR G719F expressed high level of phosphorylated EGFR without EGF induction and this is comparable to that of cells transduced with EGFR L858R. However, contradictory to our hypothesis, NIH-3T3 cells transduced with EGFR G729E expressed negligible level of phosphorylated EGFR without EGF induction (Fig.3.B). In addition, A549 cells expressed negligible level of phosphorylated EGFR without EGF induction (Fig.3.D); suggesting that EGFR in A549 is overexpressed, but not constitutively activated.

**Fig.1.**
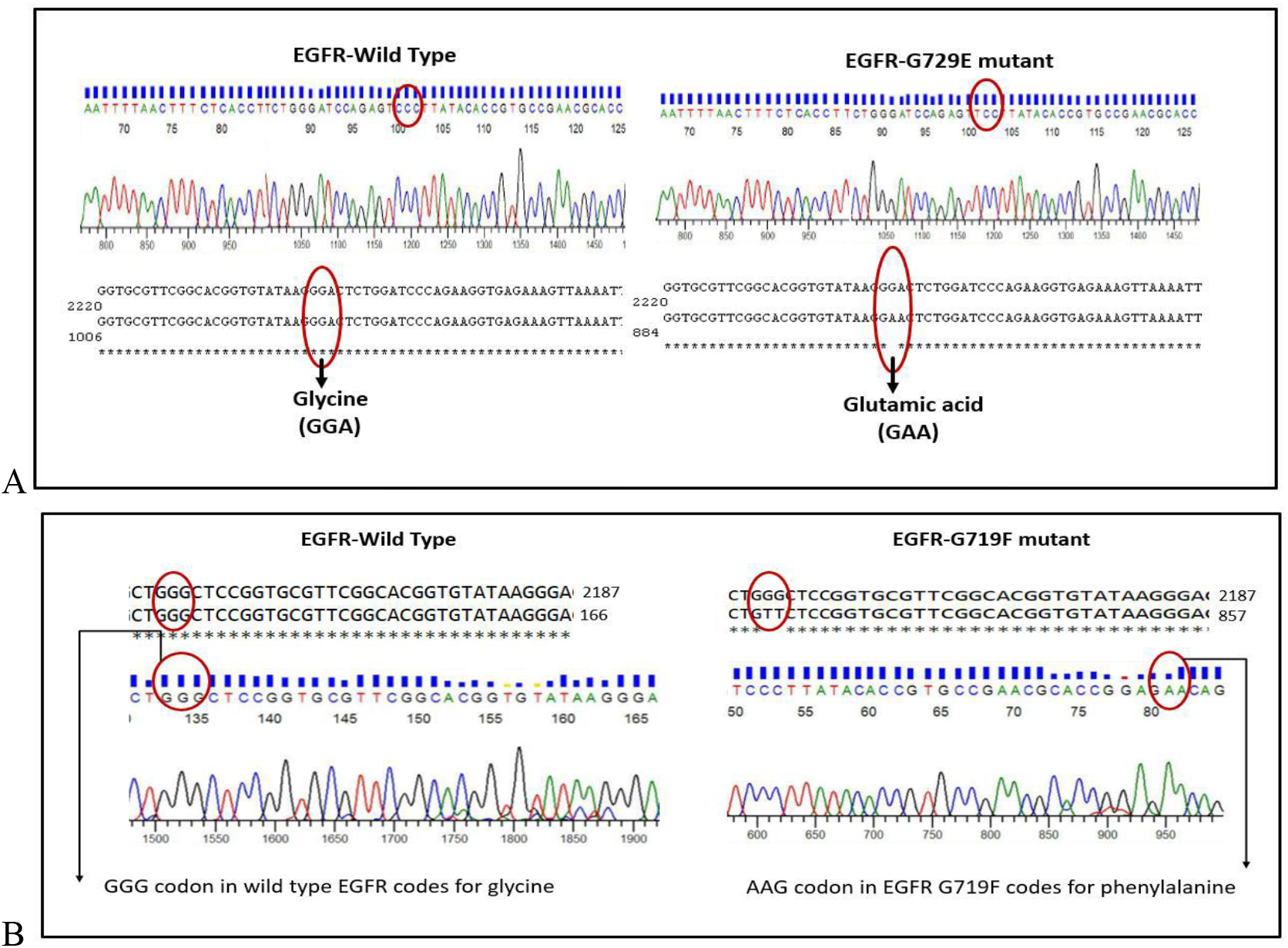
A. Clustal Omega Multiple Sequence Alignment of EGFR WT and EGFR G729E sequences. B. Clustal Omega Multiple Sequence Alignment of EGFR WT and EGFR G719F sequences.

**Fig.2.**
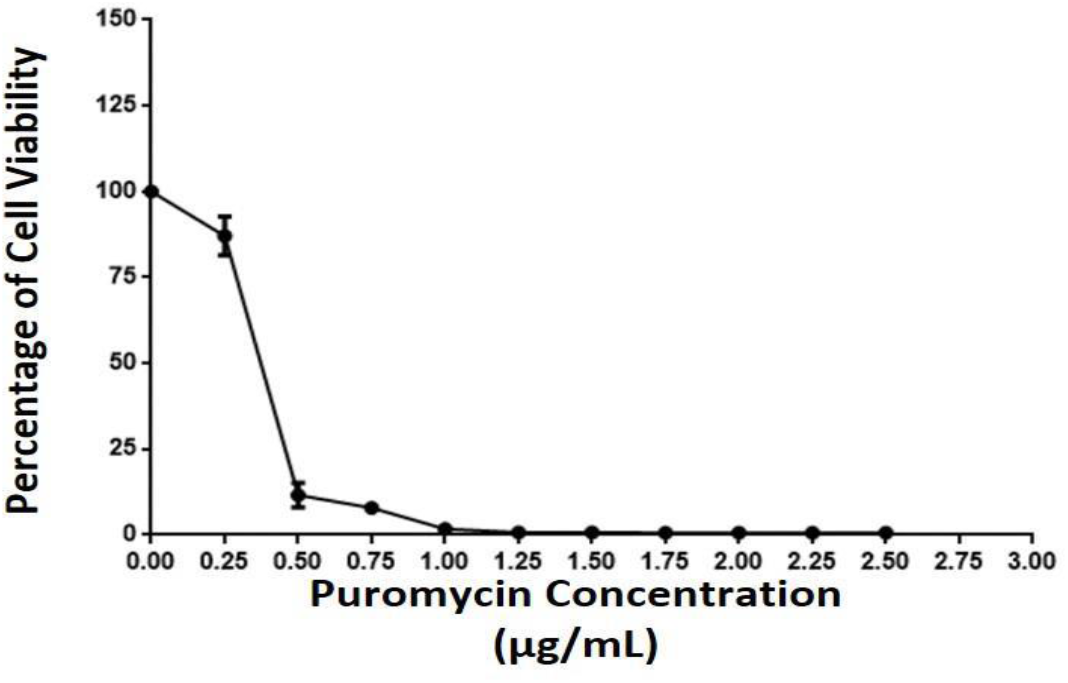
Percentage viability of NIH-3T3 cells after puromycin treatment for 7 days

**Fig.3.**
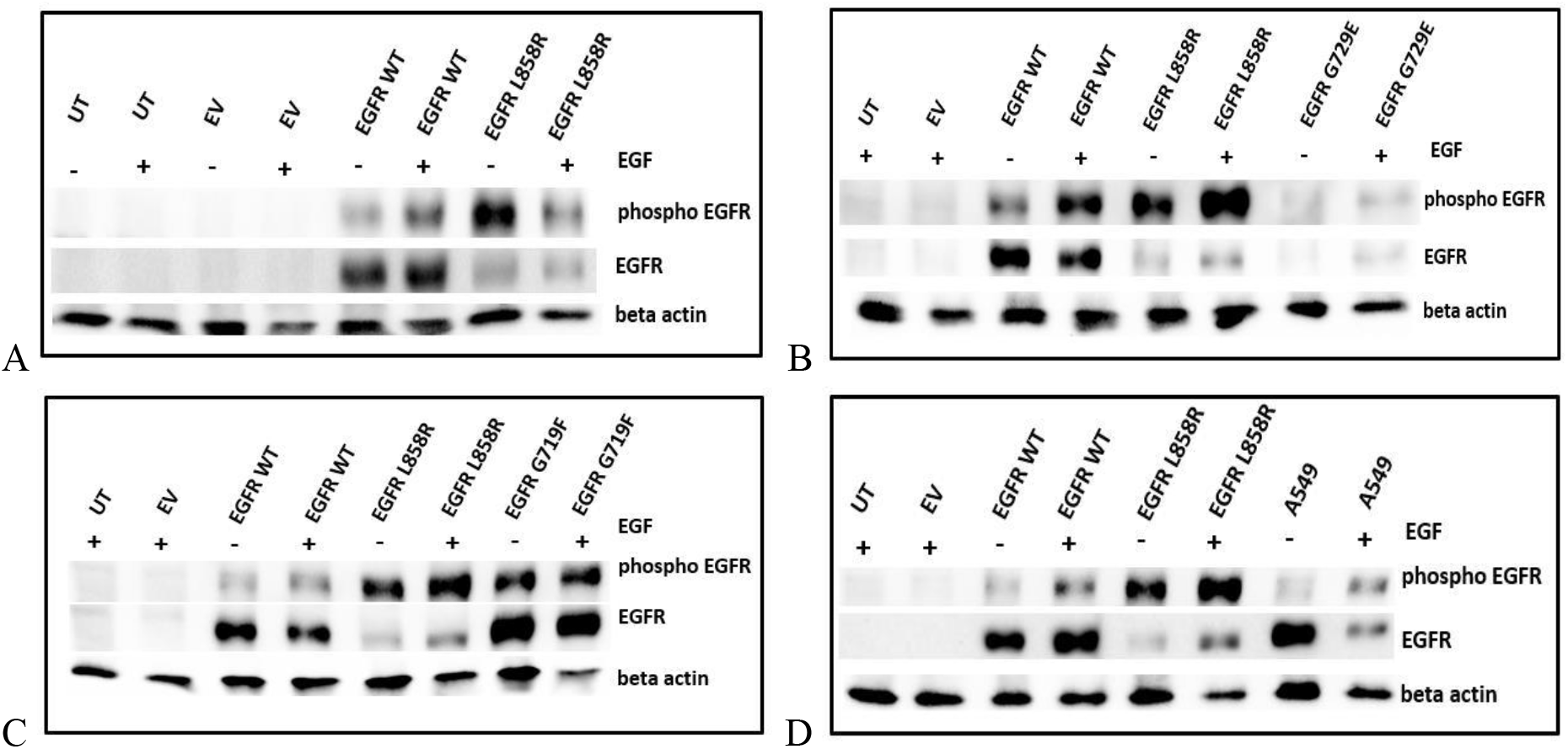
Analysis of phospho and total EGFR expression in NIH-3T3 cells without and with EGF induction: Lysates were prepared from untransduced cells (UT), cells transduced with empty vector (EV), cells transduced with wild type EGFR (EGFR WT) and cells transduced with mutant EGFR (A-EGFR L858R, B-EGFR G729E, C-EGFR G719F) or A549 cells (D)

### Analysis of ERK phosphorylation status

As observed from the Fig.4, there is negligible change in the expression levels of phosphorylated ERK among untransduced cells, cells transduced with empty vector, cells transduced with EGFR WT and cells transduced with mutant EGFR L858R/EGFR G729E/EGFR G719F or A549 cells.

**Fig.4.**
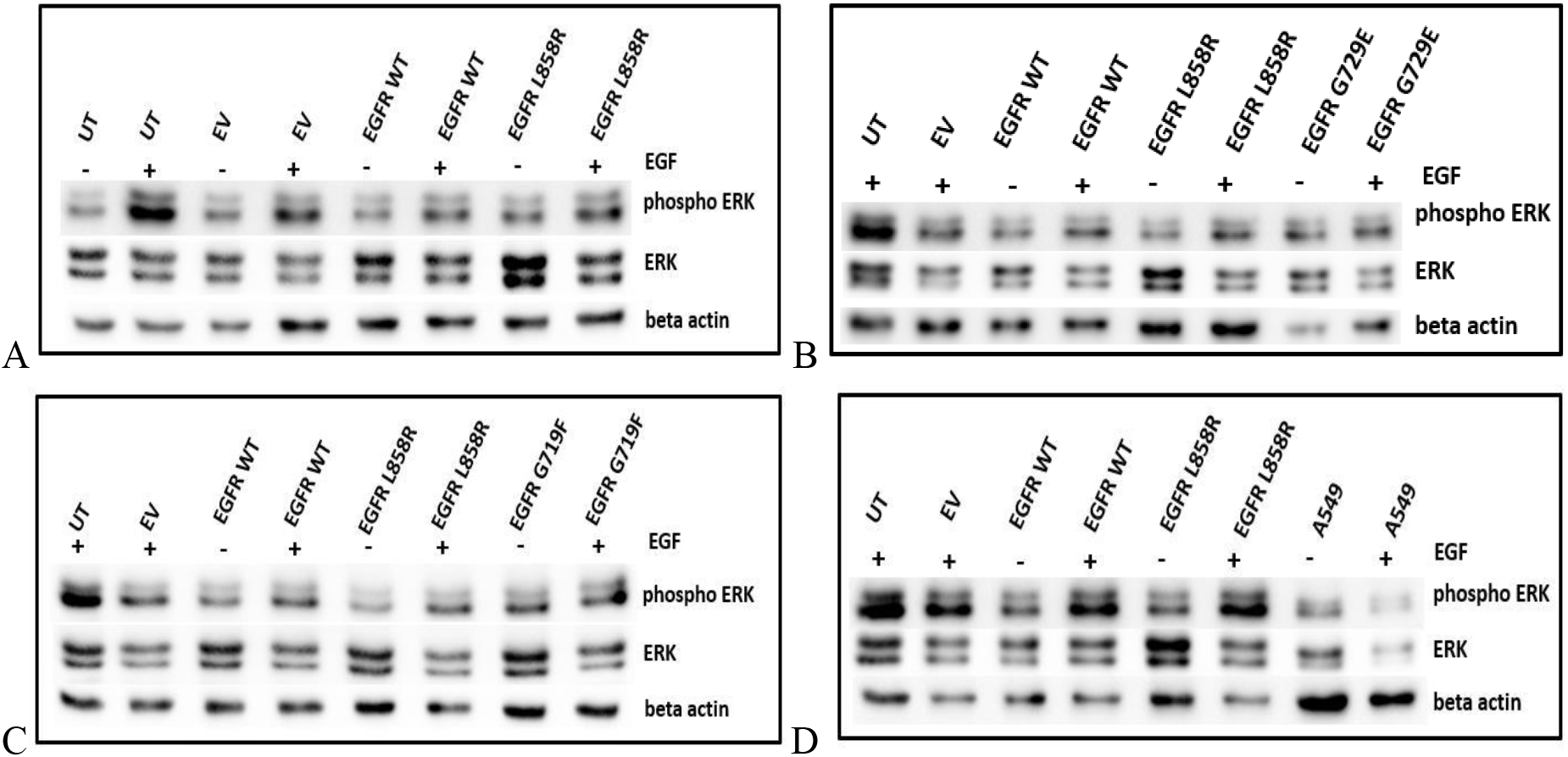
Analysis of phospho and total ERK expression in NIH-3T3 cells without and with EGF induction: Lysates were prepared from untransduced cells (UT), cells transduced with empty vector (EV), cells transduced with wild type EGFR (EGFR WT) and cells transduced with mutant EGFR (A-EGFR L858R, B-EGFR G729E, C-EGFR G719F) or A549 cells (D)

### Analysis of anchorage independent growth on soft agar

**Fig.5.**
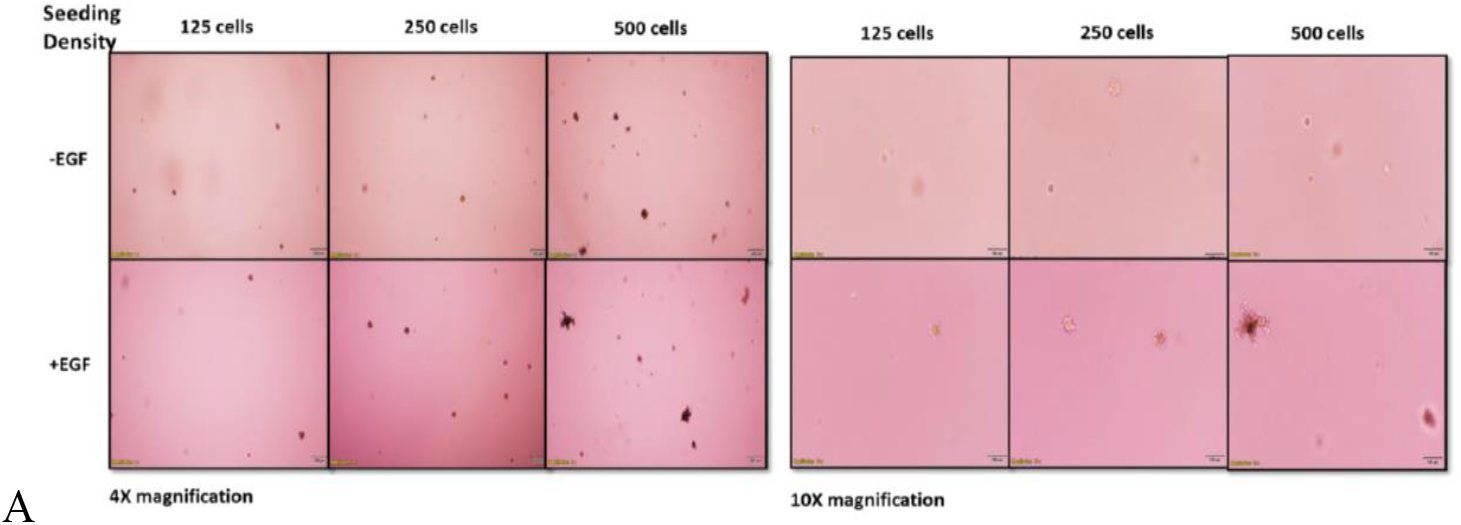
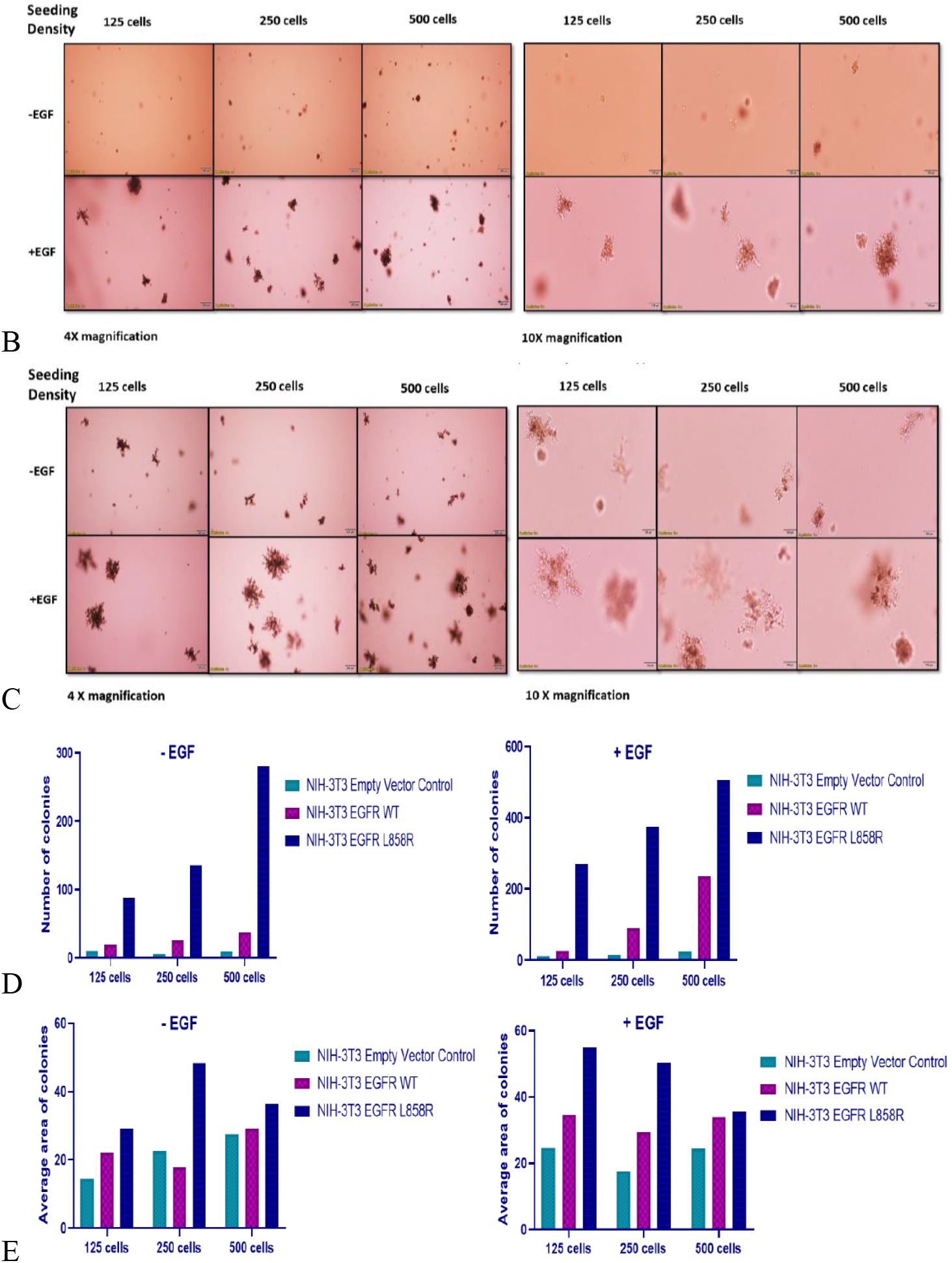
Representative images of anchorage-independent growth of NIH-3T3 cells expressing pBABE-puro empty vector (A), pBABE-puro wild type EGFR (B) and pBABE-puro EGFR L858R (C). Colony counts and average area (pixel^2^) of colonies formed by NIH-3T3 cells expressing pBABE-puro empty vector/EGFR WT/EGFR L858R are given in Figs. D and E respectively.

The principle underlying soft agar anchorage independent growth assay is that cells with oncogenic mutations can grow as colonies on an agar substratum as compared to the normal cells. Here, we showed that the colonies formed by NIH-3T3 cells expressing EGFR L858R mutation were more in number and bigger in size compared to the colonies formed by the cells expressing pBABE-puro empty vector or EGFR wild type vector (Figs A, B and C). The colony counts and average area (pixel^2^) of colonies formed by NIH-3T3 cells (Figs D and E) also confirmed the same.

## Supporting information

supplementary figue 1

