## supplementary figue 1 for "*In Vitro* Validation of Computationally Predicted Oncogenic Driver Mutations in EGFR Tyrosine Kinase Domain"

**Supplementary Figures:**

| Residue | Position | Normalized Conservation Score | Color Grade |
|---|---|---|---|
| Glycine (G) | 719 | -1.245 | 9 |
| Glycine (G) | 729 | -0.717 | 7 |
| Leucine (L) (known driver) | 858 | -1.082 | 8 |

Fig.1. Evolutionary conservation scores of selected mutation residues: scores were generated using Consurf server analysis; color grade ranges from 1-9 (variable to conserved).
